# Intramolecular interactions enhance the potency of gallinamide A analogs against *Trypanosoma cruzi*

**DOI:** 10.1101/2021.12.22.473926

**Authors:** Elany Barbosa Da Silva, Vandna Sharma, Lilian Hernandez-Alvarez, Arthur H. Tang, Alexander Stoye, Anthony J. O’Donoghue, William H. Gerwick, Richard J. Payne, James H. McKerrow, Larissa M. Podust

**Affiliations:** Skaggs School of Pharmacy and Pharmaceutical Sciences, Center for Discovery and Innovation in Parasitic Diseases, University of California San Diego, La Jolla, California; Departamento de Física, Instituto de Biociências, Letras e Ciências Exatas, Universidade Estadual Paulista Julio de Mesquita Filho, São José do Rio Preto, São Paulo, Brazil; Center for Marine Biotechnology and Biomedicine, Scripps Institution of Oceanography. University of California San Diego, La Jolla, California; School of Chemistry, The University of Sydney, Sydney, NSW 2006, Australia; Australian Research Council Centre of Excellence for Innovations in Peptide and Protein Science

## Abstract

Gallinamide A, a metabolite of the marine cyanobacterium *Schizothrix* sp., selectively inhibits cathepsin L-like cysteine proteases. We evaluated potency of gallinamide A and 23 synthetic analogs against intracellular *Trypanosoma cruzi* amastigotes and the cysteine protease, cruzain. We determined the co-crystal structures of cruzain with gallinamide A and two synthetic analogs at ∼2Å. SAR data revealed that the N-terminal end of gallinamide A is loosely bound and weakly contributes in drug-target interactions. At the C-terminus, the intramolecular π−π stacking interactions between the aromatic substituents at P1’ and P1 restrict the bioactive conformation of the inhibitors, thus minimizing the entropic loss associated with target binding. Molecular dynamics simulations showed that in the absence of an aromatic group at P1, the substituent at P1’ interacts with tryptophan-184. The P1-P1’ interactions had no effect on anti-cruzain activity whereas anti-*T. cruzi* potency increased by ∼5-fold, likely due to an increase in solubility/permeability of the analogs.

## INTRODUCTION

The World Health Organization recognizes Chagas disease, caused by the protozoan parasite *Trypanosoma cruzi*, as one of the most prevalent neglected tropical diseases in the World. In Latin America, Chagas disease is endemic, and about 8 million people are estimated to be infected worldwide^1–4^. The global healthcare and economic burden of Chagas disease is substantial^5–7^, while the clinically available treatments are limited to two drugs, benznidazole and nifurtimox. Both drugs have been recently FDA-approved for use in children of 2 to 12 years old (benznidazole) and from birth to 18 years old (nifurtimox)^8–10^, however, neither has been approved for adults. While benznidazole and nifurtimox have significant efficacy in the acute phase, both drugs suffer from the liabilities of low efficacy in chronically infected patients and adults and may cause serious adverse reactions leading to premature termination of treatment^11^. Thus, development of new anti-Chagas drugs with improved efficacy and less toxicity is a priority.

The goal of developing safer and more efficacious drugs for the treatment of Chagas disease led to an investigation of natural secondary metabolites from plants and cyanobacteria for activity against *T. cruzi*^12–14^. Cyanobacteria produce a large variety of bioactive molecules attractive for pharmaceutical applications in various fields of anti-infective drug discovery, including antibacterial, antifungal, antiviral, and antiprotozoal agents^15–17^. Tropical filamentous marine cyanobacteria have emerged as an important source of biologically active secondary metabolites^18^. The natural product gallinamide A, isolated from marine cyanobacteria discovered at a tropical reef near Piedras Gallinas on the Caribbean coast of Panama, was first identified as an inhibitor of *Plasmodium falciparum* growth^19^. Additional studies showed that the mechanism of action for this compound was due to inhibition of the food vacuole cysteine proteases that are collectively known as falcipains^20,21^. *P. falciparum* falcipain enzymes are cathepsin L-like proteases and therefore additional studies showed that gallinamide A is also a potent and highly selective inhibitor of human cathepsin L^22^. The unique linear lipopeptide structure, and potential applicability to proteases of medical relevance, provoked interest in establishing chemical syntheses of gallinamide A and structural analogs^20,23–26^. Indeed, gallinamide A has become an attractive starting point for developing selective inhibitors targeting human cathepsin L and other Clan CA cysteine proteases in human pathogens^21,23,24,27^. Based on comparison of the protein sequence alignment and substrate specificity, cathepsin L is similar to cysteine proteases from unicellular parasites that are responsible for neglected tropical diseases, such as falcipains from *P. falciparum* and cruzain from *T. cruzi*^28–30^. The cysteine protease cruzain is a promising drug target because this enzyme is essential in *T. cruzi* at all stages of the life cycle^31–33^.

Previously, fifteen gallinamide A analogs were synthesized and tested against cruzain and human cathepsin L^23^, while thirty-five additional analogs were synthesized and tested against falcipain-2 and -3 and *P. falciparum*.^21,27^ Recently, both sets of analogs were evaluated against SARS-CoV-2 and analogs **19** and **23** demonstrated efficacy against viral infection in VeroE6 cells by inhibiting host cathepsin L in the picomolar range^24^. Analog numbering in this work is according to Ashhurst et al.^24^. The gallinamide A analogs that exhibited activity against human cathepsin L also showed potent inhibition of cruzain and intracellular *T. cruzi* amastigotes, making the gallinamide scaffold a promising lead for anti-Chagas drug development^23^. Specifically, the parent gallinamide A natural product inhibited cruzain with IC50 of 0.26 nM and intracellular *T. cruzi* amastigotes with EC50 of 15 nM^23^.

In the absence of an experimental structure of a gallinamide-cathepsin L complex, molecular docking in human cathepsin L was instrumental in predicting a putative gallinamide A binding pose and guiding analog design for effective inhibition of the human protease^22,23^. Here we report the first X-ray structures of the cruzain complexes with gallinamide A and analogs that will be instrumental for developing this class of cysteine protease inhibitors as anti-Chagas agents and for other biomedical applications. We also evaluated two series of synthetic gallinamide A analogs (23 compounds) against cruzain and intracellular *T. cruzi* amastigotes. The SAR data combined with the molecular dynamics (MD) simulations, including MM-GBSA free energy decomposition and thermodynamic integrations (TI), identified hot spots and structural determinates underlying the selective inhibition of the Clan CA cysteine proteases by gallinamide A and analogs. This work lays the foundation for the design of natural product analogs with improved activities and drug-like properties.

## RESULTS

### X-ray structure analysis of the cruzain-gallinamide A complex

#### Cruzain-gallinamide A interactions

The cruzain-gallinamide A co-crystal structure containing six cruzain molecules in an asymmetric unit (chains A-F) was determined to a resolution of 2.2 Å. Data collection and refinement statistics are provided in **Table 1**. In all chains, electron density was present in the active side cleft, suggesting that gallinamide A was bound in an extended linear conformation via a covalent bond to the C25 thiol (**Fig. 1A, B**). Gallinamide A spans the S1 ’, S1, S2, and S3 pockets of the cruzain active site (**Fig. 1C, D**). Binding of gallinamide A occurs via H-bonding, aromatic stacking interactions and hydrophobic interactions. Three H-bonds are formed between the inhibitor backbone and cruzain (**Fig. 1E**). The amide NH atoms of P1 and P2 H-bond to carbonyl oxygen atoms of D161 and G66, respectively, while the carbonyl oxygen of P1’ H-bonds to the NH of the Q19 side chain.

**Table 1.** Data collection and refinement statistics

| **Data collection** |  |  |  |
| --- | --- | --- | --- |
| Inhibitor (ID) | Gallinamide A (GN9) | **23** (83K) | **24** (83E) |
| PDB ID | 7JUJ | 7S19 | 7S18 |
| Space group | P 43 21 2 | P 32 2 1 | I 41 2 2 |
| Cell dimensions |  |  |  |
| *a*, *b*, *c* (Å) | 139.85, 139.85, 163.15 | 95.93, 95.93, 68.85 | 99.50, 99.50, 85.37 |
| $\alpha$, $\beta$, $\gamma$ (°) | 90.0, 90.0, 90.0 | 90.00, 90.00, 120.00 | 90.00, 90.00, 90.00 |
| Molecules in AU | 6 | 1 | 1 |
| Wavelength | 1.11587 | 1.11587 | 1.11587 |
| Resolution (Å) | 2.20 | 2.08 | 2.14 |
| *R*<sub>sym</sub> (%) | 33.0 (882.6) | 42.0 (555.8) | 6.8 (557.3) |
| *I* / $\sigma$ *I* | 11.83 (0.43) | 8.37 (0.54) | 21.31 (0.89) |
| Completeness (%) | 99.9 (99.4) | 96.6 (83.08) | 99.9 (70.36) |
| Redundancy | 26.6 (26.8) | 19.7 (16.6) | 25.2 (26.7) |
| **Crystallization conditions** | 0.8 M K-phosphate, pH 9.7; 0.01 M betaine hydrochloride | 0.8 M K-phosphate, pH 8.0; 0.1 M CaCl<sub>2</sub>, 0.01 M betaine hydrochloride | 3.2 M NaCl, 0.1 M sodium citrate, pH 5.3; 0.01 M sarcosine. |
| **Refinement statistics** |  |  |  |
| No. reflections | 78166 | 20511 | 12135 |
| *R*<sub>work</sub> / *R*<sub>free</sub> (%) | 20.1/26.7 | 20.0/ 24.7 | 19.7/ 28.9 |
| No. atoms |  |  |  |
| Protein | 9552 | 1592 | 1589 |
| Inhibitor | 252 | 57 | 60 |
| Solvent | 233 | 79 | 3 |
| Wilson plot B | 57.3 | 37.1 | 74.3 |
| Mean B value | 64.6 | 39.3 | 87.4 |
| B-factors |  |  |  |
| Protein | 64.9 | 39.5 | 88.6 |
| Inhibitor | 91.4 | 46.5 | 94.3 |
| Solvent | 53.7 | 39.8 | 75.2 |
| R.m.s deviations |  |  |  |
| Bond lengths (Å) | 0.013 | 0.019 | 0.013 |
| Bond angles (°) | 1.816 | 2.173 | 1.898 |
| **Ramachandran statistics** |  |  |  |
| Preferred (%) | 95.0 | 95.8 | 92.5 |
| Allowed (%) | 5.0 | 3.8 | 6.6 |
| Outliers (%) | 0.0 | 0.5 | 0.9 |

<sup>1</sup>Data for the highest resolution shell are shown in parentheses.

**Figure 1.**
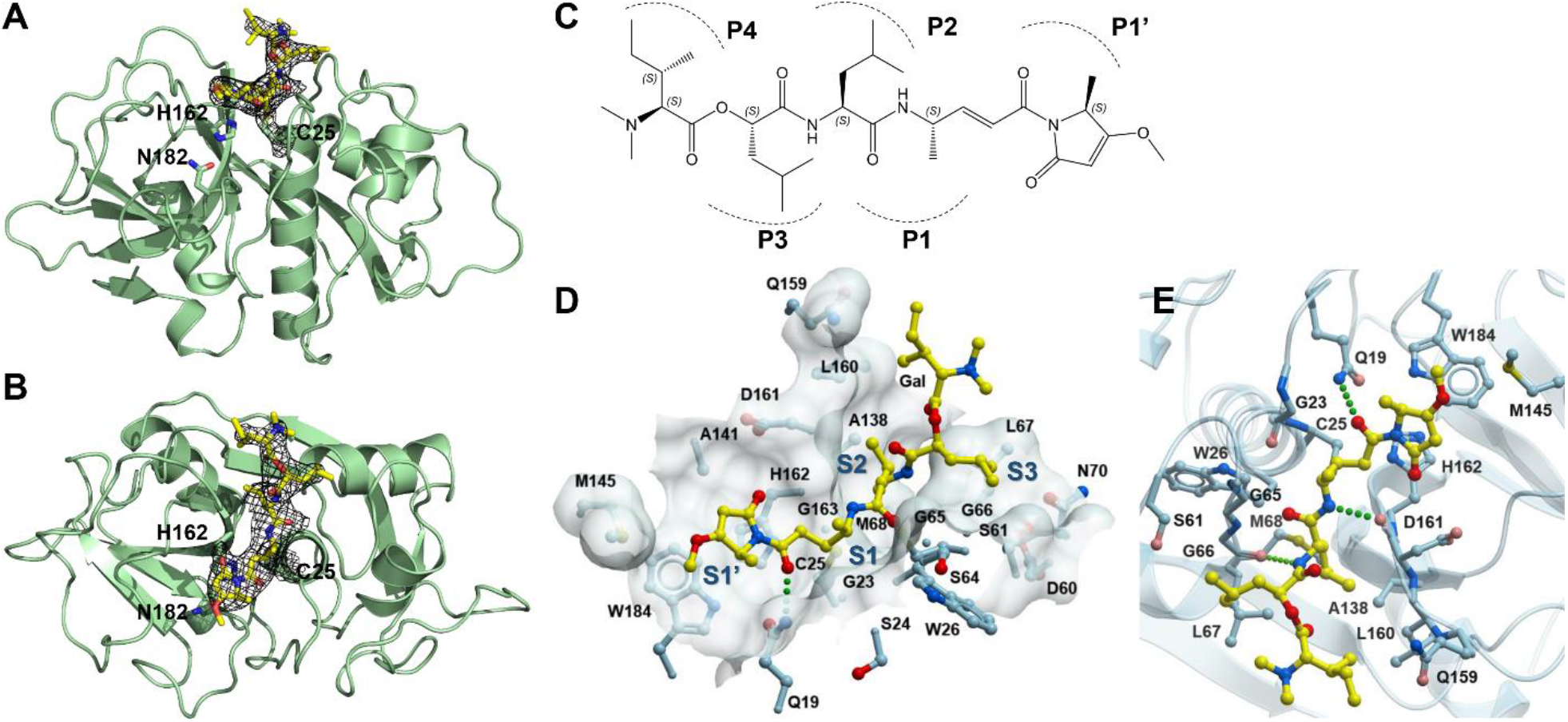
Structure of the cruzain-gallinamide A complex. **A, B**. View of the cruzain-gallinamide A complex in two orthogonal orientations. The view in B was obtained by 90° rotation of molecule in A toward the viewer. Cruzain (chain A) is shown as a green ribbon, gallinamide A is in yellow sticks, and the C25-H162-N182 acid-base-nucleophile triad is in green sticks. Electron density 2F_o_-F_c_ omit map (black mesh) is countered at 2.0 σ. **C**. Chemical structure of gallinamide A. The side chains within gallinamide A are labeled according to the Schechter−Berger nomenclature^34^, with the first, second and third residues (from left to right) labeled as P4, P3 and P2, and fourth and fifth residues labelled as P1 and P1′, respectively. **D, E**. The substrate-binding site is shown in two different orientations as a semitransparent surface (**D**) built by the amino acid residues shown in light blue and a ribbon (**E**); bound gallinamide A is in yellow balls-and-sticks. H-bonds between cruzain and gallinamide A are shown in green dots. Heteroatoms in all images are color-coded, oxygen in red, nitrogen in blue, sulfur in dark yellow.

The cyclic methylmethoxypyrrolidinone (MMP) moiety at P1’ of gallinamide A is inserted between D161 and H162 in the S1’ pocket and is stabilized via aromatic stacking interactions with W184 and H162 **(Fig. 2**). This orientation places the enone pharmacophore in close proximity to the thiol group of the acid-base-nucleophile triad C25-H162-N182 favoring formation of the irreversible covalent bond between C25 and gallinamide A via a Michael-addition-like reaction^35^. The electron density for the MMP is well defined with the exception of the 3-methoxy group due to conformational ambiguity of the latter **(Fig. 2A**). M145 in proximity of the 3-methoxy substituent of the MMP also adopts multiple conformations, suggesting a loose contact with MMP. The P2 and P3 leucine moieties of gallinamide A bind in the S2 and S3 pockets, while the *N,N*-dimethylisoleucine moiety (*N,N*-Me2-L-Ile) at P4 is exposed to the bulk solvent (**Figs. 1D, 2A**). The closest side chains to the *N,N*-Me_2_-L-Ile moiety include E207, E208, and Q159, and are located 7-8 Å away (**Fig. 2B**). Lack of interactions between the *N,N*-Me_2_-L-Ile residue and cruzain leads to binding ambiguity of the gallinamide A N-terminus and is consistent with the flat SAR at P4 position of the gallinamide analogs reported elsewhere^23^.

**Figure 2.**
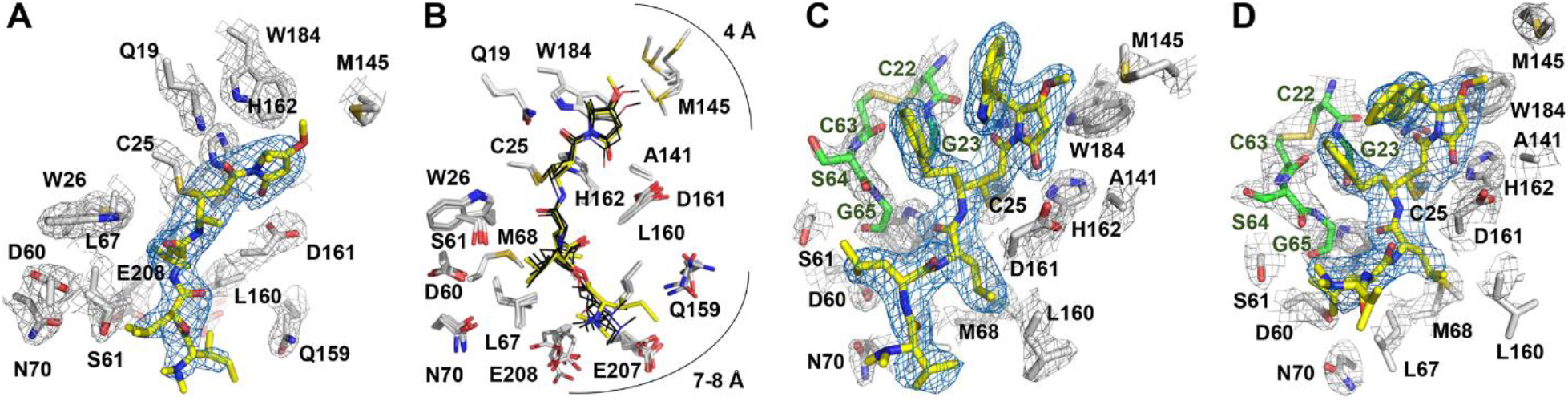
Cruzain-gallinamide A analog interactions. Fragments of the electron density 2F_o_-F_c_ omit map for gallinamide A (**A**), analog **23** (**C**) and analog **24** (**D**) are countered at 2.5 σ (blue mesh). Fragments of the electron density 2F_o_-F_c_ map for cruzain are countered at 2 σ (gray mesh). Inhibitors are highlighted in yellow, amino acid side chains are in gray, G23, C22, C63, S64 and G65 are in green; the heteroatoms are color-coded: oxygen in red, nitrogen in blue, sulfur in dark yellow. **B**. Gallinamide A binding site is shown as an overlap between the six protein chains, A-F. Gallinamide A in chain A is in yellow, in chains B-F is in black.

#### Similarity and differences between the gallinamide A docking and X-ray poses

The major differences between the computationally predicted docking (in the related cathepsin L protease) and experimental X-ray binding poses of gallinamide A is in binding of the *N,N*-Me_2_-L-Ile moiety. In the cruzain-gallinamide A X-ray structure, *N,N*-Me_2_-L-Ile is exposed to the bulk solvent instead of binding in the S3 pocket, as it was predicted by molecular docking in cathepsin L (compare cyan and yellow poses in **Fig. 3A, B**)^22,23^. While there is a possibility of having artifacts from docking, this discrepancy may also be a result of structural differences between cathepsin L and cruzain. Despite the high sequence identity (46%) and structural similarity between cathepsin L and cruzain, there are a few non-conservative amino acid substitutions in the substrate-binding site. The S3 pocket harbors three major substitutions, D60→N, S61→E and N70→Y, that distinguish the substrate binding sites of cathepsin L from cruzain (**Fig. 3C**). Three other substitutions, Q159→D, M145→L and L160→M are located in S1’ and S2 pockets. In addition, a pair of consecutive glutamate residues, E207 and E208, is absent in cathepsin L, and the equivalent positions are occupied by serine and alanine, respectively.

**Figure 3.**
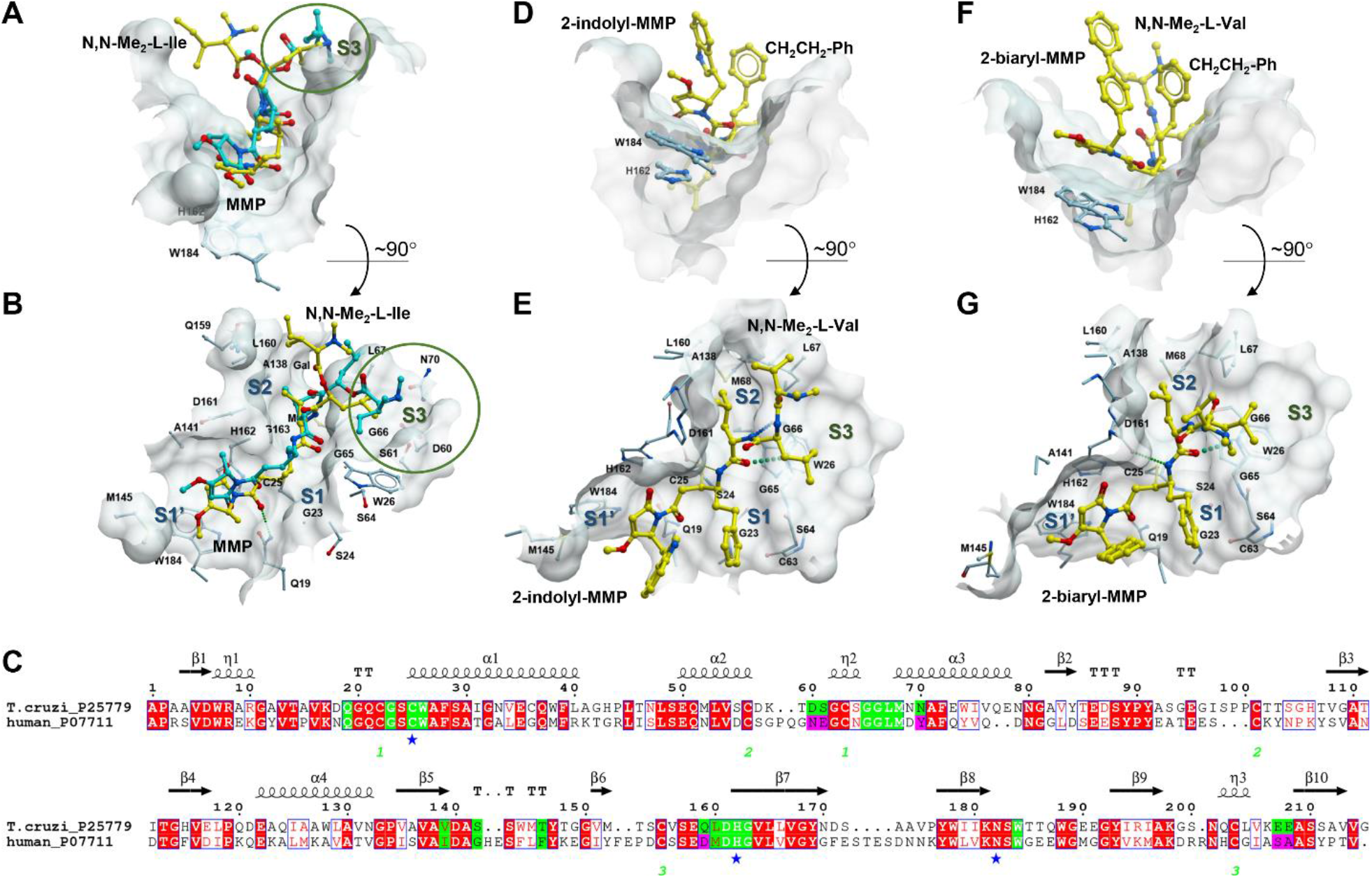
Binding modes of gallinamide A and analogs. **A, B**. Gallinamide A binding poses in cruzain (yellow) and cathepsin L (cyan) are shown in the context of the cruzain binding site (semi-transparent surface). Encircled in green is the S3 pocket harboring non-conservative amino acid substitutions D60→N, S61→E and N70→Y. **C**. Sequence alignments between cruzain and human cathepsin L. Residues constituting the substrate-binding site are in green; non-conservative substitutions between cruzain and cathepsin L are highlighted in magenta. Blue stars indicate the catalytic acid-base-nucleophile triad C25-H162-N182. Six cysteine residues involved in formation of the disulfide bonds are numbered in green. Residue numbering is according to cruzain. **D, E**. 2-indolyl-MMP analog **23** binding in cruzain. **F, G**. 2-biaryl analog **24** binding in cruzain. Inhibitors carbon atoms are in yellow, oxygen in red, nitrogen in blue.

### Biological activity of the gallinamide A analogs

#### Activity of gallinamide A analogs against cruzain and T. cruzi amastigotes

A total of 23 gallinamide A analogs from two series were evaluated against both *T. cruzi* cruzain and intracellular *T. cruzi* amastigotes (**Tables 2, 3**). The majority of the analogs screened exhibited inhibitory activity against cruzain with IC_50_ values in the low nanomolar to picomolar range with twenty analogs being more potent than the parent gallinamide A natural product. In the *T. cruzi* intracellular amastigote assay, seven analogs were equipotent to gallinamide A’a inhibitory activity, while others possessed inferior activity to the natural product, presumably due to differences in cell permeability. Finally, the LD_50_ toward the murine host cells was >0.6 µM for all tested compounds, yielding a selectivity index (SI) of 6,000 for the most potent analogs.

**Table 2.** Structure−activity relationships of gallinamide A 2-indolyl-MMP analogs **11**-**23**

| Analog | R | R<sub>1</sub> | R<sub>2</sub> | R<sub>3</sub> | Cruzain IC<sub>50</sub> (nM)<sup>a</sup> | *T. cruzi* EC<sub>50</sub> (nM)<sup>b</sup> |
| --- | --- | --- | --- | --- | --- | --- |
| Gallinamide A | *N,N*-Me<sub>2</sub>-L-Ile | CH<sub>3</sub> | L-leucine | L-leucine | 1.3 ± 0.1 | 14.7 ± 2.3 |
| 11 | *N,N*-Me<sub>2</sub>-L-Val | CH<sub>3</sub> | L-leucine | L-leucine | 0.11 ± 0.0025 | 85 ± 25 |
| 12 | *N,N*-Me<sub>2</sub>-L-Val | CH<sub>3</sub> | CH<sub>2</sub>-cyclohexyl | CH<sub>2</sub>-cyclohexyl | 0.74 ± 0.095 | 30 ± 5 |
| 13 | *N,N*-Me<sub>2</sub>-L-Val | CH<sub>3</sub> | CH<sub>2</sub>-cyclohexyl | L-leucine | 0.34 ± 0.11 | 58 ± 19 |
| 14 | *N,N*-Me<sub>2</sub>-L-Val | CH<sub>3</sub> | L-leucine | CH<sub>2</sub>-cyclohexyl | 0.081 ± 0.013 | 12 ± 0.15 |
| 15 | *N,N*-Me<sub>2</sub>-L-Val | CH<sub>3</sub> | CH<sub>2</sub>CH<sub>2</sub>-Ph | L-leucine | 0.49 ± 0.089 | >600 |
| 16 | 4-(*N*-Me)piperidine | CH<sub>3</sub> | CH<sub>2</sub>-cyclohexyl | L-leucine | 1.35 ± 0.15 | >600 |
| 17 | 4-(*N*-Me)piperidine | CH<sub>3</sub> | L-leucine | CH<sub>2</sub>-cyclohexyl | 0.086 ± 0.0061 | >600 |
| 18 | 4-(*N*-Me)piperidine | CH<sub>3</sub> | CH<sub>2</sub>-cyclohexyl | CH<sub>2</sub>-cyclohexyl | 4.95 ± 1.25 | 196 ± 5 |
| 19 | *N,N*-Me<sub>2</sub>-L-Val | CH<sub>3</sub> | CH<sub>2</sub>CH<sub>2</sub>-Ph | CH<sub>2</sub>-cyclohexyl | 1.14 ± 0.04 | 40 ± 3 |
| 20 | 4-(*N*-Me)piperidine | CH<sub>3</sub> | L-leucine | L-leucine | 0.19 ± 0.02 | >600 |
| 21 | *N,N*-Me<sub>2</sub>-L-Val | CH<sub>3</sub> | L-leucine | CH<sub>2</sub>-Ph | 0.85 ± 0.07 | 56 ± 4 |
| 22 | *N,N*-Me<sub>2</sub>-L-Val | Cyclohexyl | L-leucine | CH<sub>2</sub>-cyclohexyl | 0.71 ± 0.03 | 18 ± 6 |
| 23 | *N,N*-Me<sub>2</sub>-L-Val | CH<sub>2</sub>CH<sub>2</sub>-Ph | L-leucine | L-leucine | 0.09 ± 0.0091 | 18 ± 3 |
| E-64 | - | - | - | - | 29.73 ± 9.16 | - |
| Benznidazole | - | - | - | - | - | 1910 ± 700 |

<sup>a</sup>IC<sub>50</sub> values represent the average of two independent experiments performed in triplicate with at least 8 compound concentrations. Errors are given by the ratio between the standard deviation and the square root of the number of measurements.
<sup>b</sup>EC<sub>50</sub> are the average of two independent biological replicates performed in duplicate, +/- standard error of the mean. The LD<sub>50</sub> to the murine host cells was >0.6 μM for all tested compounds.

**Table 3.** Structure−activity relationships of gallinamide A 2-biaryl-MMP analogs **24**-**33**.

**Analogs 24-33**
| Analog | R | R<sub>1</sub>' | Cruzain IC<sub>50</sub> (nM)<sup>a</sup> | *T. cruzi* EC<sub>50</sub> (nM)<sup>b</sup> |
| --- | --- | --- | --- | --- |
| **Gallinamide A** | *N,N*-Me<sub>2</sub>-L-Ile | CH<sub>3</sub> | 1.3 ± 0.1 | 14.7 ± 2.3 |
| **24** | *N,N*-Me<sub>2</sub>-L-Val | Ph | 0.12 ± 0.01 | 13 ± 2 |
| **25** | 4-(*N*-Me)piperidine | Ph | 0.40 ± 0.1 | 222 ± 2 |
| **26** | *N,N*-Me<sub>2</sub>-L-Val | 4-CF<sub>3</sub>Ph | 0.68 ± 0.1 | 78 ± 8 |
| **27** | 4-(*N*-Me)piperidine | 4-CF<sub>3</sub>Ph | 0.79 ± 0.1 | 420 ± 61 |
| **28** | *N,N*-Me<sub>2</sub>-L-Val | 4-OCH<sub>3</sub>Ph | 0.28 ± 0.01 | 19 ± 4 |
| **29** | 4-(*N*-Me)piperidine | 4-OCH<sub>3</sub>Ph | 0.26 ± 0.04 | 261 ± 90 |
| **30** | *N,N*-Me<sub>2</sub>-L-Val | 4-CNPh | 0.11 ± 0.01 | 19 ± 7 |
| **31** | 4-(*N*-Me)piperidine | 4-CNPh | 0.13 ± 0.009 | 423 ± 100 |
| **32** | *N,N*-Me<sub>2</sub>-L-Val | Naph | 0.69 ± 0.03 | 59 ± 30 |
| **33** | 4-(*N*-Me)piperidine | Naph | 1.56 ± 0.7 | 521 ± 173 |
| **E-64** | - | - | 29.73 ± 9.16 | - |
| **Benznidazole** | - | - | - | 1910 ± 700 |

<sup>a</sup>IC<sub>50</sub> values represent the average of two independent experiments which were determined based on at least 8 compound concentrations in triplicate. Errors are given by the ratio between the standard deviation and the square root of the number of measurements.
<sup>b</sup>EC<sub>50</sub> are the average of two independent biological replicates in duplicate, +/- standard error of the mean. The LD<sub>50</sub> to the murine host cells was >0.6 μM for all tested compounds.

The first analog series assessed was composed of the thirteen 2-indolyl-MMP analogs **11**-**23** (**Table 2**) featuring three amino acid modules within the linear chain plus two different P4 functionalities, 4-*N*-methylpiperidine (4-(*N*-Me)piperidine) or *N,N*-Me_2_-L-Val, at the pseudo-N-terminus. At the C-terminal end, an indole functionality (derived from L-tryptophan) was appended to the MMP ring via a methylene bridge. All substituents at P1, P2 and P3 were introduced on the common 2-indolyl-MMP scaffold. A majority (85%) of the compounds in this series showed higher activity against cruzain than gallinamide A, with **14, 17**, and **23** being >15-fold more potent. Two analogs, **14** and **23**, having N-terminal R=*N,N*-Me_2_-L-Val retained potency against *T. cruzi* amastigotes, while **17** and other analogs (**16, 18** and **20**) having more hydrophobic R=4-(*N*-Me)piperidine at the N-terminus lost potency against *T. cruzi*. Importantly, the potent activity against the cruzain target by the analogs carrying the 4-(*N*-Me)piperidine substituent (**16, 17, 18** and **20**) is consistent with exposure of the N-terminal moiety observed in the crystal structures.

The hydrophobic CH_2_-cyclohexyl substituent was explored at P2 and P3 positions alone and in combination with the L-leucine side chain (compounds **11-14**). CH_2_-cyclohexyl was tolerated in both positions with the most potent configuration achieved in **14**, where R_2_=L-leucine and R_3_=CH_2_-cyclohexyl. A bulky aromatic substituent CH_2_CH_2_-Ph was introduced in **23** at P1 where it replaced L-alanine of the parental scaffold. In **23**, R_1_=CH_2_CH_2_-Ph was combined with R_2_=R_3_=L-leucine that did not affect activity against the cruzain target (compared to **11**) but enhanced approximately 5-fold the potency of **23** against *T. cruzi* amastigotes. Interestingly, the same CH_2_CH_2_-Ph group at P2 or P3 positions reduced activity against *T. cruzi* amastigotes (compounds **15, 19** and **21**). A substitution pattern with R_2_=CH_2_CH_2_-Ph and R_3_=L-leucine largely eliminated activity of **15** against *T. cruzi*.

Introduction of biaryl-substituted MMP moieties at the C-terminus was explored by evaluating analogs **24**-**33** (**Table 3**). These compounds possessed either 4-(*N*-Me)piperidine or *N,N*-Me_2_-L-Val at the pseudo-N-terminus and CH_2_CH_2_-Ph at the P1 site^24,27^. All but one of the analogs of this series exhibited more potent activity against cruzain compared to gallinamide A. However, only three analogs, **24, 28** and **30**, retained the anti-*T. cruzi* activity. Similar to the 2-indolyl-MMP series, analogs carrying R=4-(*N*-Me)piperidine (compounds **25, 27, 29, 31** and **33**) had anti-parasitic activity inferior to the counterparts carrying R=*N,N*-Me2-L-Val. Furthermore, analogs having more hydrophobic R1’=4-CF_3_-Ph and R1’=Naph experienced an approximately 4-5-fold decrease in anti-*T. cruzi* activity.

### X-ray structure analysis of the cruzain-23 and cruzain-24 complexes

We also determined the co-crystal structures of cruzain covalently bound to gallinamide A analogs **23** from the 2-indolyl-MMP series (**Table 2**) and **24** from the 2-biaryl-MMP series (**Table 3**). Both analogs are among the most potent cruzain and *T. cruzi* inhibitors from this collection. Unlike cruzain-gallinamide A, the co-crystal structure of each analog contained one molecule in the asymmetric unit. Electron density for analogs was well defined with the exception of the exposed N-terminal *N,N*-dimethylvaline moiety (*N,N*-Me_2_-L-Val) (**Fig. 2C, D**), particularly in **24** where it pointed away from the cruzain surface (**Fig. 3D-G**). Binding ambiguity of *N,N*-Me_2_-L-Val resembles that of *N,N*-Me_2_-L-Ile in the parent gallinamide A natural product. The P2 and P3 leucine moieties interact in the S2 and S3 pockets, respectively (**Fig. 3D-G**).

Interestingly, the indolyl moiety of **23** and the biaryl moiety of **24** established intramolecular π-π stacking interactions with the P1 substituent, CH_2_CH_2_-Ph in the molecule (**Fig. 3D, F**). While CH_2_CH_2_-Ph made surface contacts with the backbone atoms of G23, C22, C63, S64 and G65 (shown in green in **Fig. 2C, D**), neither indolyl nor biaryl moieties interacted with the protein. Intramolecular interactions of the CH_2_CH_2_-Ph at P1 with the indolyl or biaryl moieties at P1’ may reduce the hydrophobic surface, increase solubility in aqueous medium, and positively impact permeability and potency of **23** and **24**. Furthermore, these intramolecular interactions may also stabilize the biologically active conformation and reduce the entropic loss associated with the binding to the target. We speculate that the role of the aromatic MMP substituents is responsible for restricting the flexibility of the analogs to lock them into their biologically active conformation. Given that no amino acid side chains are involved, these intramolecular interactions are expected to broadly enhance activity against Clan CA cysteine proteases. The **23** and **24** co-crystal structures were determined in different space groups that minimize any impact of crystal packing interactions on the inhibitor binding poses.

### Molecular modeling

#### Preferential binding modes of **14, 17** and **23** revealed by MD simulations and per-residue free energy decomposition

To explore dynamics of cruzain-ligand interactions *in silico*, we performed MD simulations for the top hits of the 2-indolyl-MMP series, compounds **14, 17** and **23**, for which cruzain inhibitory activity exceeded 15-fold that of gallinamide A. A critical structural difference that distinguishes **14** and **17** from **23** is the absence of the CH_2_CH_2_-Ph moiety at P1 that excludes the stabilizing effect of the intramolecular interactions between the indolyl and CH_2_CH_2_-Ph moieties in **14** and **17**. We also calculated the per-residue free energy decomposition (Δ*G*_*res*_) employing the structure ensemble generated for each complex simulation. As shown in **Fig. 4**, residues W184, L160 and L67 established non-polar interactions with the 2-indolyl-MMP, L-leucine, CH_2_-cyclohexyl and *N,N*-Me_2_-L-Val moieties at P1’, P2, P3 and P4 positions, respectively. In resemblance to the conformation adopted by the L-leucine at P3 of gallinamide A, the CH_2_-cyclohexyl moiety of **14** and **17** project towards and interact with L67. The MD simulations showed that residues Q19, G66 and D161 are establishing H-bonds with the backbone of the gallinamide A analogs, which is in agreement with the interactions observed from the X-ray crystal structures^36–38^.

**Figure 4.**
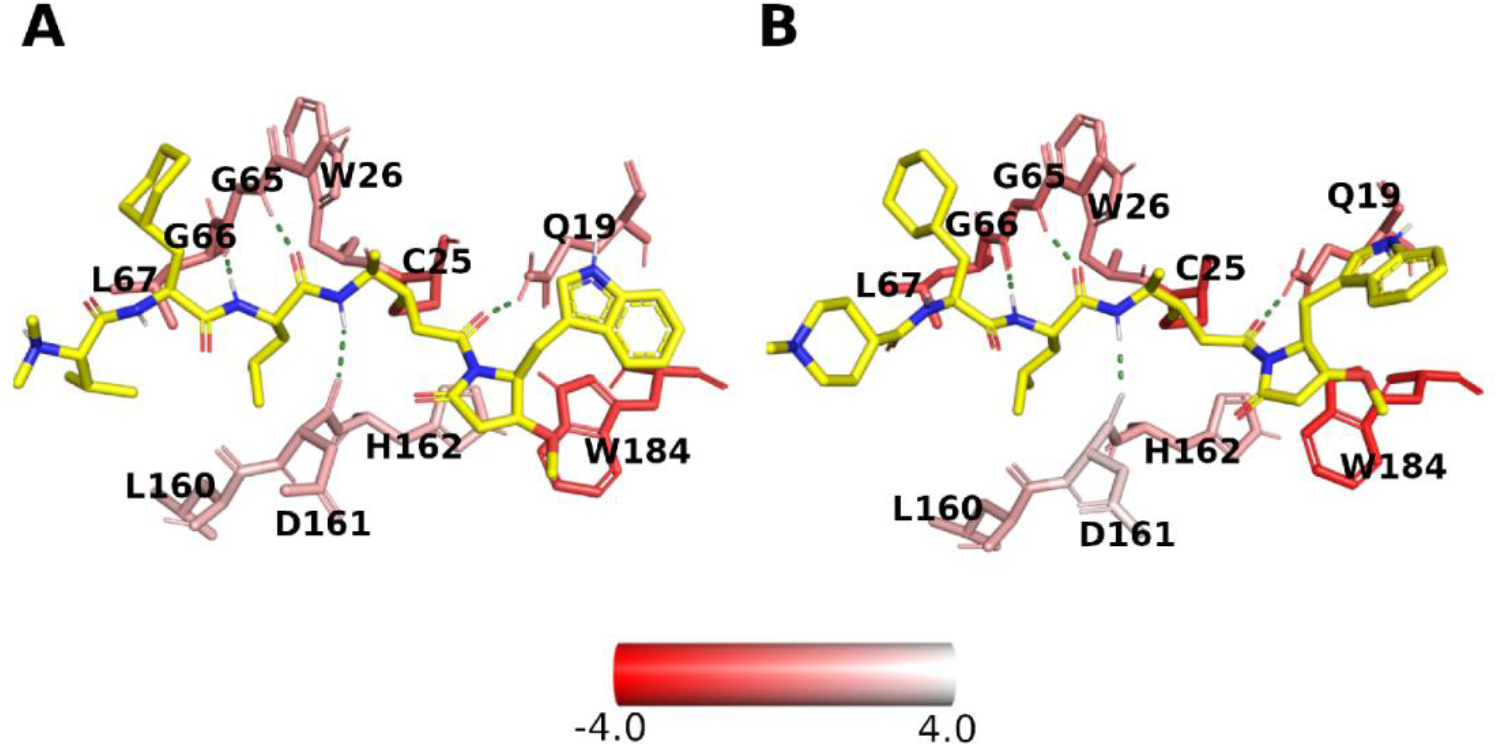
Representative structures calculated from the MD simulations of cruzain bound to 14 (A) and 17 (B). **14** and **17** are shown in yellow sticks. The cruzain binding site residues are depicted as sticks and colored according to the per-residue energy contributions (ΔG_res_). The ΔG_res_ values are expressed in kcal/mol. Hydrogen bonds are displayed as green-dashed lines, and the main interacting residues are labeled.

As the electron density corresponding to the P4 moiety of gallinamide A and the analogs was not well-defined (**Fig. 3**), we computationally assessed the dynamics of the P4 fragment along the course of the MD simulations. Note that compounds **14** and **23** have the *N,N*-Me_2_-L-Val group at P4 site while compound **17** contains the 4-(*N*-Me)piperidine group at this position. To describe the changes in the orientation of the P4 moiety within the cruzain binding site, we monitored over time the dihedral angle ψ (between the equivalent C and CA atoms) of the P4-P3 linkage calculated for each of the three complexes. We observed that the cyclohexyl ring at P3 established more stable interactions with S3 when the P4 position was occupied by *N,N*-Me_2_-L-Val (**Fig. 5A**). In contrast, with 4-(*N*-Me)piperidine at P4, two well-defined populations of the cyclohexyl ring at P3 sampled by ψ angle were observed. One population bound within S3 while the other one was exposed to the solvent (**Fig. 5B**). These results suggest that the 4-(*N*-Me)piperidine group placed in the P4 position may destabilize interactions of cyclohexyl moiety at S3.

**Figure 5.**
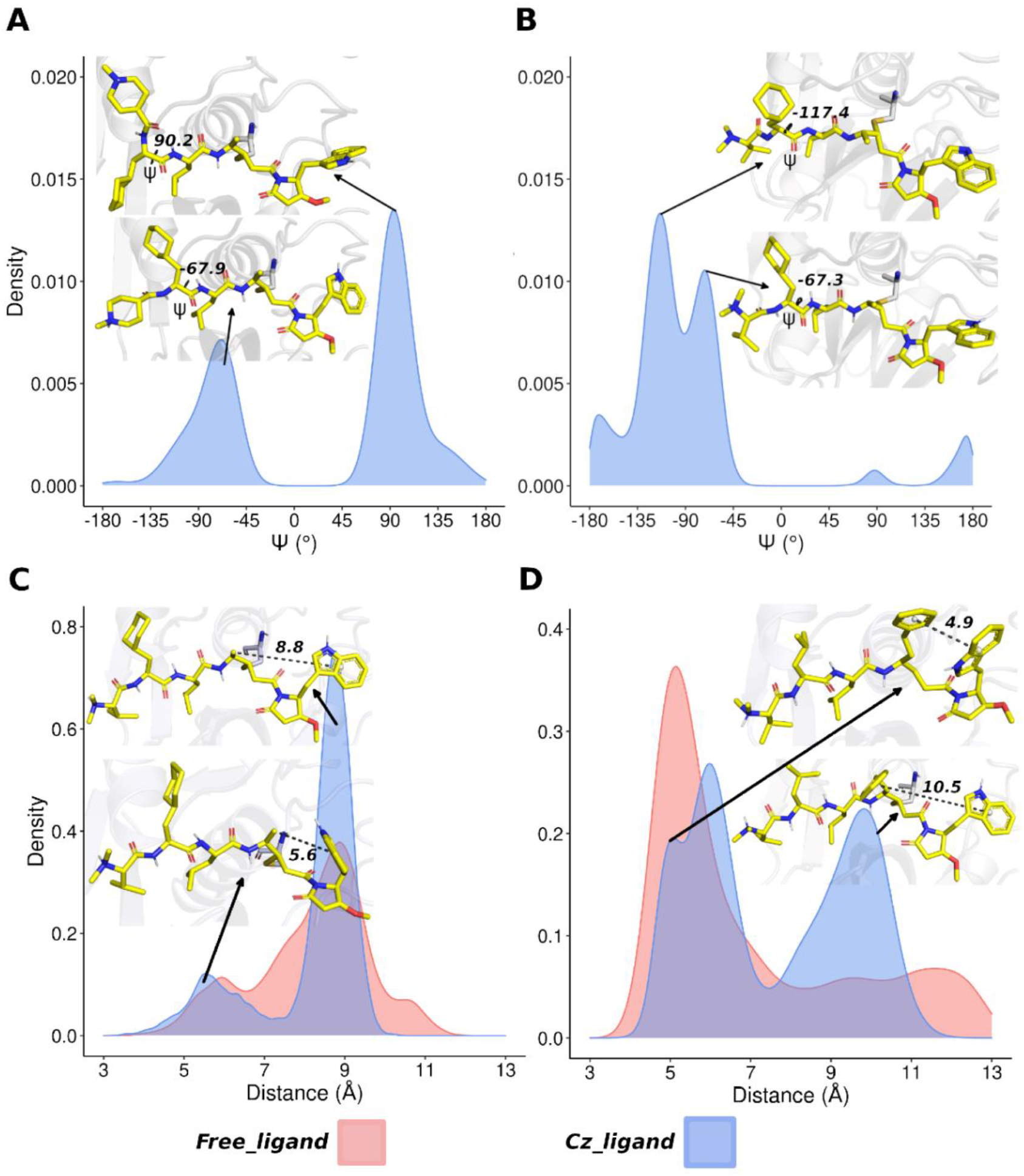
Binding conformations of 14, 17 and 23 revealed by MD simulations. Ψ values of the P3 moiety sampled along the MD simulation trajectory are plotted for cruzain-bound **14** (**A**) and **17** (**B**). Distribution of distances between the center of mass of phenyl and indole rings along the MD simulation trajectory are plotted for **14** (**C**) and **23** (**D**), cruzain-bound (blue) and free in a water box (red). The representative structure of each ψ angle and each distance population is depicted above the respective peak. Gallinamide A analogs are shown in yellow sticks. The ψ angles and distances are represented by dashed lines labelled with the corresponding values.

The MD simulations showed that the predicted interactions of the indolyl substituent in **14** and **17** differ from those observed in the co-crystal structure of the **23-**cruzain complex (compare **Fig. 4A** and **4B** with **Fig. 2C**). In the absence of the CH_2_CH_2_-Ph group at P1, the indole ring of **14** and **17** mainly interacts with W184 (**Fig. 5C**) that is consistent with the critical role of W184 in the interactions with these analogs (see energy values of W184 in **Fig. 4**). Molecular dynamic simulations of the **23**-cruzain complex showed that the indole group of **23** may occur in both conformations: (1) interacting with W184 and (2) establishing intramolecular π−π stacking interactions with the CH_2_CH_2_-Ph moiety at P1 (**Fig. 5D**). When free in solution, **23** is mostly forming intramolecular π−π stacking interactions as observed in the co-crystal structure of the **23-**cruzain complex (**Fig. 5D**). Given that **14** and **23** have the same activity against cruzain (**Table 2**), this might suggest that the gain in binding enthalpy due to intermolecular interactions of the indole ring in **14** is comparable to the entropy loss associated with the intramolecular interactions of the indole ring in **23**.

#### Prediction of relative binding affinities of the gallinamide A analogs for cruzain by thermodynamic integration free energy calculations

Because in each experimentally tested gallinamide A derivative more than one moiety was substituted, we computationally assessed the contribution of each individual modification for the affinity for cruzain. For this purpose, we performed ΔΔ*G*_*calc*_ calculations of cruzain-gallinamide A analogs relative to the cruzain-gallinamide A complex, employing TI free energy calculations. The thermodynamic cycle involving the alchemical transformations of the inhibitor free in solution and covalently bound to the enzyme is shown in **Fig. 6**. The results of computation are shown in **Table 4**, where positive values indicate that the native complex possesses higher affinity. We conclude that the individual inclusion of CH_2_-cyclohexyl at R_3_, and CH_2_CH_2_-Ph at R_2_ significantly enhanced affinity for cruzain. Addition of the indole ring at R’ also increased the compound affinity, albeit not as much as the aforementioned modifications. Conversely, 4-(*N*-Me)piperidine at P4 resulted in a weaker binding compared to the *N,N*-Me_2_-L-Ile present in gallinamide A. Finally, the incorporation of *N,N*-Me_2_-L-Val at P4 does not seem to change the affinity of the derivative. These results are in agreement with the X-ray structures showing ambiguous *N,N*-Me_2_-L-Val binding (**Fig. 3C, D**) and experimental IC_50_ values shown in **Tables 2 and 3**. This suggests that the per-residue free energy decomposition computational approach could be very useful for evaluating new chemical modifications prior to chemical synthesis of the analogs.

**Figure 6.**
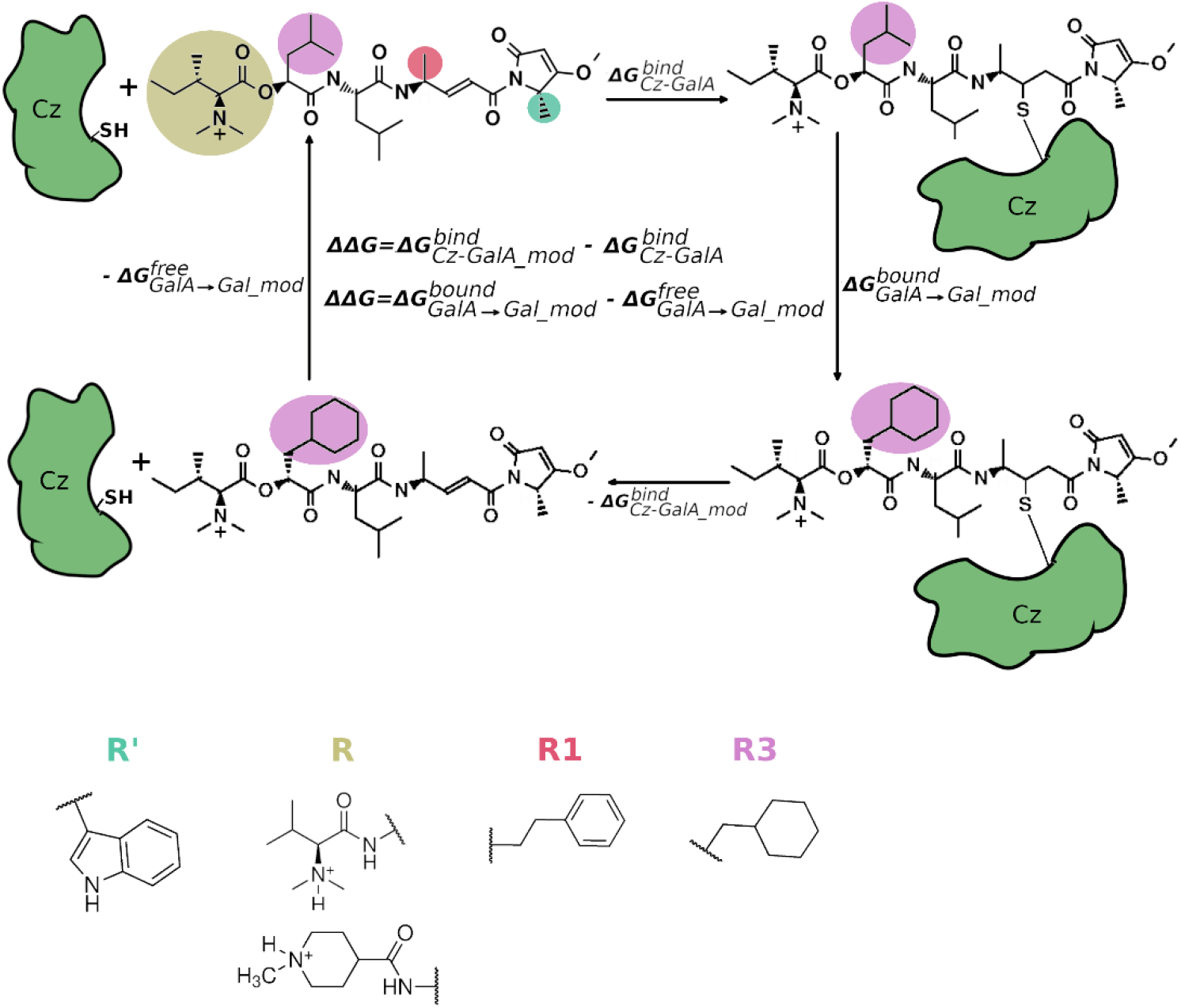
Thermodynamic cycles used to calculate the ΔΔG_calc_ values for the covalent complexes. Thermodynamic cycle involves the gallinamide A and analogs free in solution and covalently-bonded to cruzain (in green). Each moiety that was chemically-modified within the gallinamide A structure is colored differentially; corresponding chemical modifications are shown below the cycle diagram. The ΔΔG_calc_ values were calculated according to the equation depicted inside the cycle. NB: Cz = *T. cruzi* cruzain.

**Table 4.** Per-residue free energies differences between the complexes of cruzain-gallinamide A and cruzain-gallinamide A analogs

| Chemically-modified group | $\Delta\Delta G_{\text{Coulomb}}^a$<br>(kcal/mol) | $\Delta\Delta G_{\text{vdw}}^a$<br>(kcal/mol) | $\Delta\Delta G_{\text{calc}}^{a,b}$<br>(kcal/mol) |
| --- | --- | --- | --- |
| Indolyl pyrrolidinone at P' | $0.65 \pm 0.11^c$ | $-2.79 \pm 0.08$ | $-2.13 \pm 0.14$ |
| CH<sub>2</sub>-cyclohexyl at P3 | $-2.36 \pm 0.09$ | $-0.85 \pm 0.17$ | $-3.21 \pm 0.19$ |
| CH<sub>2</sub>CH<sub>2</sub>-Ph at P2 | $-2.76 \pm 0.07$ | $-0.28 \pm 0.09$ | $-3.04 \pm 0.11$ |
| 4-(N-Me)piperidine at P4 | $1.31 \pm 0.14$ | $0.39 \pm 0.11$ | $1.70 \pm 0.18$ |
| N,N-Me<sub>2</sub>-L-Val at P4 | $-1.03 \pm 0.08$ | $0.86 \pm 0.14$ | $-0.17 \pm 0.16$ |

<sup>a</sup> $\Delta\Delta G_X = \Delta G_X(\text{gallinamide A}) - \Delta G_X(\text{derivative})$ , where X stands for Coulomb (charge-related transformations), vdw (van der Waals transformations), and calc (calculated relative free energy).
<sup>b</sup> $\Delta\Delta G_{\text{calc}} = \Delta\Delta G_{\text{Coulomb}} + \Delta\Delta G_{\text{vdw}}$
<sup>c</sup>Final standard error of mean calculated by error propagation

## DISCUSSION

The inhibitory potential of the indolyl-substituted MMP moiety in gallinamide A analogs was first recognized in the studies on the falcipain proteases from *P. falciparum*. The most potent *P. falciparum* falcipain-2 and -3 inhibitor (IC_50_^falcipain-2^ = 12.0 nM, IC_50_^falcipain-3^ = 66.7 nM M, EC_50_^*P. falciparum*^ = 9.7 nM) carried an 2-indolyl-MMP, similar to the analogs **11**-**23**^21^. One of these analogs was able to cure murine malaria infection in mice at 25 mg/kg and delayed parasitemia at ≥6.25 mg/kg^27^. We show in this work that gallinamide A and a range of synthetic analogs are also potent inhibitors of *T. cruzi* cruzain (IC_50_ of 1 nM) and intracellular *T. cruzi* amastigotes (EC_50_ of 15 nM)^23^.

The co-crystal structures of the gallinamide drug–target complexes determined as part of this work will aid in hit-to-lead optimization of this class of protease inhibitors to improve metabolic stability, solubility and permeability, while maintaining high potency and selectivity. The structures demonstrated that the N-terminal P4 moiety makes loose contacts with cruzain and may be used to improve solubility and permeability of the next generation of the anti-Chagas gallinamide A analogs. Exposure of the P4 moiety in cruzain contrasts to its proposed binding in the S3 pocket of cathepsin L as predicted by molecular docking^23^. The discrepancy between the docking and crystallographic poses may be a docking artifact or the result of non-conservative amino acid substitutions in S3 of cathepsin L affecting pocket shape and electronic properties. Determination of the experimental cathepsin L-gallinamide A structure would be instrumental to resolving this ambiguity and may also have additional applications for the development of SARS-CoV-2 entry inhibitors^24^.

Activity data for the P2 and P3 substituents (**Tables 2, 3**) is consistent with earlier studies by others aimed at the development of cruzain inhibitors. Specifically, bulky groups at P3, such as carboxybenzyl or 3-pyridinyl, are present in known potent irreversible cruzain inhibitors^39–41^. Similarly, the CH_2_-cyclohexyl group at P3 is well tolerated in the gallinamide analogs. The L-phenylalanine and to a lesser extent L-leucine and L-valine are commonly present at P2^42,43^. These groups were embedded within potent gallinamide A analogs used in this study.

SAR trends observed for the P1 site suggest a preference for a center with S-configuration^40,44,45^. This preference was also observed when gallinamide A analogs were tested against cruzain and cathepsin L. D-Phe at the P1 site results in ≥100-fold reduction in activity^23^. Inhibitors with a homophenylalanine at P1, which tends to be more resistant to metabolism, showed high potency^40^. The β-branched alkyl substituents, particularly L-Val, were also beneficial for activity^44^.

The most striking observation made in this work is that CH_2_CH_2_-Ph at P1 is involved in intramolecular π−π stacking interactions with the aromatic indolyl or biaryl substituents of MMP at P1’. Neither indolyl nor biaryl moieties contact the protein target, suggesting that in the context of **23** and **24** their role is in restricting the flexible inhibitor into its bioactive conformation and minimizing the entropic loss associated with binding to the target. The conformational restriction (rigidification) of a flexible ligand is a commonly used strategy in drug design that enhances potency, improves selectivity and reduces drug metabolism^46^. The same rigidification principle, either via non-covalent intramolecular interactions or via introducing a covalent linkage, may be explored in designing peptidomimetic inhibitors targeting cathepsin L-like proteases. The gallinamide A and analogs co-crystal structures determined in this work will also have utility in assessing the contributions of the individual substituents for the binding affinity of gallinamide A analogs to Clan CA cysteine proteases. Combining this structural data with the per-residue free energy decomposition approach provides the possibility of validation of new chemical modifications computationally prior to chemical synthesis of future gallinamide A analogs.

## EXPERIMENTAL SECTION

### Chemicals

Gallinamide A and its analogs were synthesized and characterized previously by the HPLC analysis to be >95%.^24,27^ Analog numbering is according to^24^. Dulbecco’s modified Eagle medium was from Thermo Scientific (Paisley, UK); penicillin−streptomycin from Omega Scientific (Tarzana, USA), fetal bovine serum from Sigma (St. Louis, USA) and benznidazole (*N*-benzyl-2-nitro-1*H*-imidazole-1-acetamide) were from Sigma (St. Louis, USA).

### Zymogen procruzain expression and activation

The pET21a vector (Novagen/EMD) containing the gene sequence encoding the C-terminally truncated His6-tagged zymogen procruzain (Δc; GenBank entry M84342.1) was used for protein expression. Procruzain was expressed in ArcticExpress (DE3) RIL cells (Agilent) and purified as described by Silva *et al* ^47^. Procruzain was activated at 0.1 M sodium acetate, pH 5.3, 10 mM EDTA, 0.9 M NaCl, and 5 mM dithiothreitol (DTT). Then DTT was removed using NAP-5 desalting columns (GE Healthcare) and the activation buffer without reducing agents. Covalent reversible inhibitor, S-methyl methanethiosulfonate (MMTS) or gallinamide A, was added to protect the catalytic thiol group of C25 and prevent self-degradation of cruzain. The cruzain-gallinamide A complex was directly used for crystallography studies. Covalently inhibited cruzain tolerated a short-term storage at -80 °C. The detailed purification and activation protocols are provided in Supplementary Information S1.

Modifications to the protocol were made to obtain the cruzain-gallinamide A analog complexes. Following procruzain activation as described above, DTT was removed using HiPrep 26/10 desalting column (GE Healthcare) saturated with the activation buffer with no DTT and supplemented with MMTS. Eluted protein fractions were immediately tested for activity. Fractions containing the MMTS-cruzain complex were immediately pooled, concentrated to ∼3 mg/mL using Amicon Ultra centrifugal filter (3 kDa molecular weight cut-off) (Millipore) and loaded onto the size-exclusion Superdex 75 10/300 GL (GE Healthcare) column saturated with 2 mM Bis-Tris, pH 5.8. To prevent cruzain autodegradation, excess MMTS was added to each fraction immediately upon elution from the column. Homogenously purified MMTS-cruzain complex was further concentrated to 5 mg/mL and stored at -80 °C for inhibitor exchange and co-crystallization.

### Enzymatic activity assay

Cruzain activity was measured by monitoring the cleavage of the fluorescent substrate, Z-Phe-Arg-aminomethylcoumarin (Z-FR-AMC), using a Synergy HTX (Biotek) fluorimeter. A 10 mM stock solution of Z-FR-AMC was prepared in dimethyl sulfoxide (DMSO). All assays were performed in black flat-bottom 96-well plates (Costar, catalog 3915), in 200 µl of 0.1 M sodium acetate, pH 5.5, supplemented with 1 mM DTT, 0.01% Triton X-100, 0.5 nM cruzain, and 2.5 µM of Z-FR-AMC ^48^. Prior to addition of the substrate, enzyme was incubated with the test compounds for 10 minutes. Following the substrate addition, fluorescent signal was recorded. Enzymatic activity was calculated from the initial rates of the reaction. Two independent experiments were performed in triplicate. Half-maximal inhibitory concentration (IC_50_) was determined by nonlinear regression analysis of the initial velocity vs. inhibitor concentration plot using Prism ^49^. At least eight inhibitor concentrations were used to build each curve. DMSO and trans-Epoxysuccinyl-L-leucylamido(4-guanidino)butane (E64) were used as negative and positive controls, respectively.

### Crystallization, data collection and structure determination

For cruzain-gallinamide A complex, gallinamide A-bound cruzain was concentrated to 10 mg/mL and the activation buffer was exchanged to 2 mM Bis-Tris pH 5.8 using a centrifugal filter with molecular weight cut-off of 3 kDa (Millipore). For cruzain-gallinamide A analog complexes, 1.2 molar excess of the analog in DMSO was added to cruzain-MMTS complex followed by adding 10-molar excess of DTT. The reaction mix was incubated for 60 minutes at room temperature for inhibitor exchange. Screening of crystallization conditions was performed using commercial high-throughput screening kits available in deep-well format from Hampton Research (Aliso Viejo, CA) or Qiagen (Germantown, MD), a nanoliter drop-setting Mosquito robot (TTP LabTech, Melbourn, UK) operating with 96-well plates, and a hanging drop crystallization protocol. For diffraction quality, crystals were further optimized in 96-well plates configured using the Dragonfly robot (TTP LabTech, Melbourn, UK) and the Designer software (TTP LabTech, Melbourn, UK). Crystals were obtained at 23 °C from the conditions specified in **Table 1**.

Diffraction data were collected remotely at beamline 8.3.1, Advanced Light Source, Lawrence Berkeley National Laboratory. Data indexing, integration, and scaling were conducted using XDS ^50^. Cruzain structure (PDB ID 3KKU)^51^ was used as a molecular replacement model. The PHASER and REFMAC5 modules of the CCP4 software^52^ suite were used to solve and refine the structure. COOT software^53^ was used for the model building. Data collection and refinement statistics are shown in **Table 1**.

### *T. cruzi* cell-based assay

Antiparasitic activity and cytotoxicity of compounds were determined as described by Boudreau *et al* ^23^. Mouse myoblasts, cell line C2C12 (ATCCCRL-1772), were maintained in Dulbecco’s modified Eagle medium (DMEM - Invitrogen) containing 4.5 g/l glucose, supplemented with 1% penicillin−streptomycin 10,000 U/mL (Invitrogen) and 5% fetal bovine serum (Sigma) at 37 °C with 5% CO_2_. *T. cruzi* CA-I/72 strain was maintained by weekly coinfection with C2C12 cells. To assess the anti-parasitic activity of the compounds, cells and parasites were seeded in 384-well black clear-bottom plates at 1 × 10^5^ parasites/mL and 2 × 10^4^ C2C12/mL density in 50 μL of DMEM media per well. Compounds were evaluated at 10 serially diluted concentrations (from 600 nM to 1.2 nM). Plates were then fixed with 4% formaldehyde for at least 2 h and stained with 0.5 μg/mL of 4′,6-diamidino-2-phenylindole (DAPI) for at least 4 hours prior to reading. Plates were imaged and analyzed with the automated ImageXpress MicroXL microscope (Molecular Devices). Infection levels (parasites per host cell) were normalized to positive control (50 µM benznidazole) and negative control (DMSO). EC_50_ (antiparasitic activity) and CC_50_ (host cell cytotoxicity) values were determined from the dose-response curves using Prism software (GraphPad Software, La Jolla, CA)^49^. Two independent assays were performed, each in duplicate.

### Parametrization of gallinamide A and analogs for computational studies

To model the covalently bound equivalents, the enone group of each compound was reduced to the *trans*-isomer. Further, the compounds were subjected to geometry optimization with single-point (SP) calculations using Gaussian 09 package^54^ with Hartree-Fock level of 163 theory, 6-31G(d) basis and Merz-Kollman (MK) scheme^55^. To obtain the atom-centered partial charges, the electrostatic potential (ESP) derived from quantum mechanical calculations were fitted using the RESP algorithm implemented in Antechamber20^56^. The compounds parameters were obtained from the Generalized Amber Force-Field 2 (GAFF2) force field^56^. Finally, the charges obtained through RESP fits underwent certain modifications, so that all equivalent atoms outside the thermodynamic integration (TI) regions across the studied compounds had the same charge, and those involved in alchemical transformations had a 0 or +1 net charge.

### MD simulations

The starting structure of cruzain-inhibitor complexes were obtained by transforming gallinamide A in the co-crystal structure determined in this work into each individual analog using Avogadro program^57^. Protonation states of cruzain residues were determined at pH=5.5 using the PDB2PQR server^58^. Systems setup was performed with tleap program from AmberTools20^56^, and a covalent bond was built between the appropriate C atom of enone moiety of each ligand to the S atom of the catalytic cysteine. AMBER14SB force field (ff14SB)^59^ was employed for protein parameters. All complexes were solvated with explicit TIP3P water molecules^60^ in an octahedral box extending at least 10 Å from the solute surface. Systems were neutralized by replacing water molecules with Na+ and Cl-counterions, depending on their net charges.

All simulations were conducted with pmemd.cuda of Amber20^56^. 50,000 energy minimization (EM) steps were performed using a combination of steepest descent and conjugate gradient procedures. The equilibration procedure was carried out in two sequential steps, i.e., the NVT and NPT ensembles. During the NVT equilibration, the heating was performed employing a linear temperature gradient from 10 to 298 K. The subsequent NPT equilibration was performed at a constant temperature of 298 K. Both equilibration steps were performed for 500 ps with the solute heavy atoms restrained with a 10.0 kcal·mol^-1^·Å^-2^ restraint constant. The Berendsen barostat and thermostat were employed in each equilibration step^61^. Finally, the non-constrained MD simulation was performed at constant pressure and temperature (1 atm and 298 K, respectively), employing the Berendsen barostat^61^ and the Langevin thermostat^62^, respectively. The particle mesh Ewald (PME) method was used to handle long-range electrostatic interactions^63^ and a distance cutoff of 10 Å was employed. The MD integration step was set to 2 fs and one snapshot was sampled every 10,000 steps. Three replicas of 200 ns were simulated in all cruzain-compound complexes by assigning different random velocities to the systems’ atoms during the respective heating steps.

### Trajectory analyses

The cpptraj program of AmberTools20 package^56^ was employed to analyze all MD trajectories and also for calculating the ψ angle (formed between the planes defined by O-C-CA-CB atoms) which describes the dynamics of P3 and P4 positions of gallinamide A analogs. Hydrogen bonds established between gallinamide A analogs and cruzain were calculated with the default geometric definition of cpptraj, i.e., a distance cutoff ≤3.0 Å between acceptor and donor atom, and the acceptor-hydrogen-donor angle ≥135°^56^.

### Per-residue free energy decomposition with Molecular Mechanics Generalized Born Surface Area (MM-GBSA)

A per-residue effective free energy decomposition was carried out in order to determine the most important residues involved in cruzain-ligand interactions^64,65^. Effective binding free energies (Δ*G*_*eff*_), which do not include the contribution of configurational entropy, were conducted for complexes of cruzain with the top hits of gallinamide A analogs using MMPBSA.py^66^. The single-trajectory approximation was used for this calculation, employing the MD simulations of aforementioned complexes. The GB-neck2 model (igb=8) with mbondi3 radii was used for estimating the polar solvation energy (Δ*G*_*GB*_)^56^. A salt concentration of 0.1 M and the default dielectric constant value (ε=1) were set. A per-residue effective free energy decomposition was carried out in order to determine the most important residues involved in cruzain-ligand interactions^64,66^. Energetically-relevant residues, i.e., hot-spots, at the interfaces of the studied complexes were predicted using the energy decomposition protocol implemented in MMPBSA.py program.

### TI Free Energy Calculations

To assess the contribution of each chemical modification to the affinity for cruzain, we perform rigorous alchemical transformation and relative free energy calculations (ΔΔ*G*_*calc*_). For this purpose, the new chemical modifications present in three best hits (compounds **14, 17**, and **23** from the series of 2-indolyl-MMP analogs) were independently analyzed. TI ΔΔ*G*_*calc*_ calculations were performed using the thermodynamic cycle reported by Hernandez-Gonzalez *et al*. for covalent ligands of cysteine proteases^67^. The resulting binding free energy of gallinamide A and analogs was derived from the sum of the various steps in which each ligand is desolvated and subsequently introduced into the cruzain binding site. The calculations were performed for each ligand in solution and within the enzyme binding site. Simulations of cruzain complexes and solvated free ligands were prepared in the same manner as the conventional MD simulations. The representative structure of each studied complex, obtained from the RMSD-clustering analysis of the equilibrium MD simulations, was selected for TI free energy calculations.

The discharge, charge, and van der Waals steps of the ligand in solution and in complex form were slowly decoupled along the simulation by changing the λ-parameter from 0 to 1 with a stride of 0.05. A total of 21 windows were used for the decoupling process. For each window, a short EM, a 1 ns NVT heating process followed by a 12 ns production run was performed in the NPT ensemble. The pressure in the NPT MD simulations was controlled by means of the Monte Carlo barostat^56^. The timestep used for all simulations was 2 fs and the PME algorithm was used to treat long-range electrostatic interactions with a cut-off of 10 Å. All MD simulations were performed using pmemd.cuda from Amber20^56^. The results were analyzed using the alchemical_analysis.py python tool^68^. Charge-related alchemical transformations involving charged moieties were conducted following a single box/dual system approach in order to keep the simulation box neutral at all λ values^69^. The protein-ligand complex and the free ligand were placed in an 73×73×78 Å cuboid box with a minimal distance of 16 Å between. Their interaction was prevented by applying a harmonic restraint (k=50 kcal·mol^-1^·Å^-2^) to the Cα atom of A30 in cruzain and the CA atom of the moiety placed at P2 position in the free ligand. Errors were propagated to calculate the total standard error of means (SEMs) of the final ΔΔ*G*_*calc*_ values.

## ASSOCIATED CONTENT

### Supporting Information

The detailed protocols for cruzain expression, purification and activation are provided in the Supporting Information S1.

### Accession Codes

Coordinates and structure factors of the cruzain-gallinamide complexes are available in the Protein Data Bank (PDB) under accession codes 7JUJ, 7S19 and 7S18. Authors will release the atomic coordinates and experimental data upon article publication.

## AUTHOR INFORMATION

### Notes

The authors declare no competing financial interest.

## ACKNOWLEDGEMENTS

We thank the staff members of beamline 8.3.1, James Holton, George Meigs and Kathryn Burnett, at the Advanced Light Source at Lawrence Berkeley National Laboratory, for assistance with data collection; Jair Lage Siqueira-Neto for access to the BSL2 instrumentation and facilities.

## ABBBREVIATIONS USED

Benznidazole: *N*-benzyl-2-nitro-1*H*-imidazole-1-acetamide
DAPI: 4’,6-diamidino-2-phenylindole
DMEM: Dulbecco’s modified Eagle medium
DMSO: dimethyl sulfoxide
DTT: dithiothreitol
EM: energy minimization
ESP: electrostatic potential
E-64: *trans*-epoxysuccinyl-L-leucylamido(4-guanidino)butane
ff14SB: AMBER14SB force field
GAFF2: Generalized Amber Force-Field 2
IC_50_: half-maximal inhibitory concentration
MD: molecular dynamics
MK: Merz-Kollman
MM-GBSA: Molecular Mechanics Generalized Born Surface Area
MMP: methylmethoxypyrrolidinone
MMTS: S-methyl methanethiosulfonate
4-(*N*-Me)piperidine: 4-(*N*-methyl)piperidine
*N,N*-Me_2_-L-Ile: *N,N*-dimethylisoleucine
*N,N*-Me_2_-L-Val: *N,N*-dimethylvaline
PDB: Protein Data Bank
PME: particle mesh Ewald
SEMs: standard error of means
SI: selectivity index
SP: single-point
TI: thermodynamic integrations
ΔΔ*G*_*calc*_: relative free energy calculations
Δ*G*_*eff*_: effective binding free energies
Δ*G*_*GB*_: polar solvation energy
Δ*G*_*res*_: per-residue free energy decomposition
Z-FR-AMC: Z-Phe-Arg-aminomethylcoumarin

## Table of Contents graphic

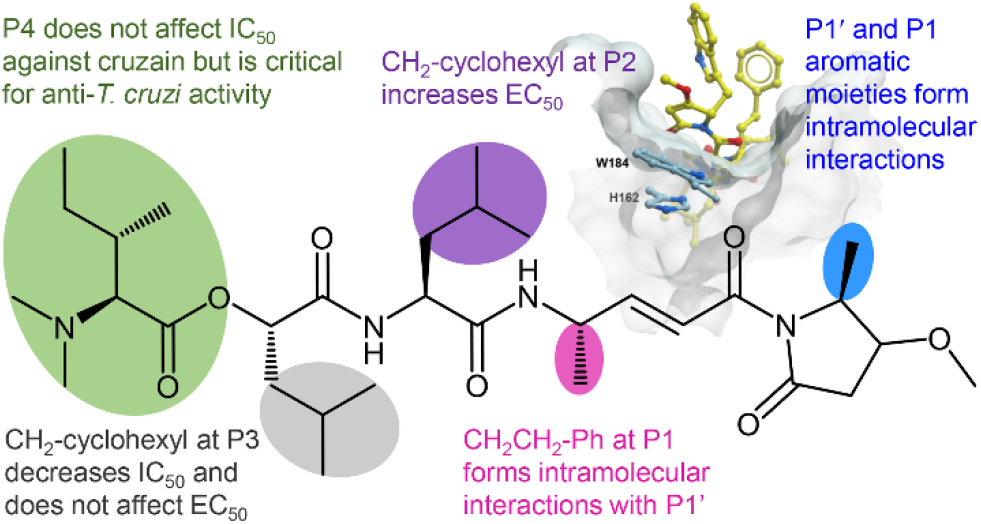

