## Supplementary Information S1 for "Intramolecular interactions enhance the potency of gallinamide A analogs against *Trypanosoma cruzi*"

**This PDF file includes:**

**Pages S2-S3:** Supporting Information S1 – Detailed protocols for cruzain expression, purification and activation.

**Zymogen procruzain expression.** The pET21a vector (Novagen/EMD) containing the gene sequence encoding the C-terminally truncated 6xHis-tagged zymogen procruzain (Δc; GenBank entry M84342.1) was used for protein expression. The procruzain was expressed in ArcticExpress (DE3) RIL cells (Agilent) and purified as described by Silva et al [47]. Recombinant cruzain was expressed and purified using a modified version of the method described by Silva *et al* [47]. Starter cultures used to express His6-tagged zymogen procruzain consisted of 5 mL of Luria broth containing ampicillin (100 μg/mL) and gentamycin (20 μg/mL). After being shaken for >15 h at 37 °C, the overnight LB starter culture was added directly to 1 L of autoinduction medium N-5052 (1% N-Z-amine, 0.5% yeast extract, 25 mM $Na_2HPO_4$, 25 mM $KH_2PO_4$, 50 mM $NH_4Cl$, 5 mM $Na_2SO_4$, 2 mM $MgSO_4$, 0.5% glycerol, 0.05% glucose and 0.2% lactose)[70] containing ampicillin (100 μg/mL) and gentamycin (20 μg/mL) in baffled 2 L shaker flasks. After incubation for 48 h, cells were collected by centrifugation (6000 rpm, 10 minutes, 4 °C). Cell pellet was suspended in 50 mL of lysis buffer per liter of expression medium and lysed under pressure in microfluidizer. The lysis buffer consisted of 50 mM Tris pH 10, 300 mM NaCl, 10 mM imidazole, 1 mM $CaCl_2$, 1 mM $MgSO_4$, a dash of DNase I (Sigma), 2 μM phenylmethanesulfonyl fluoride (PMSF), and 2 mM S-methyl methanethiosulfonate (MMTS). The lysate was clarified by centrifugation (20,000 rpm, 1 h, 4 °C).

**Zymogen procruzain purification.** All purification steps were carried out at 4°C. Recombinant procruzain was purified from the supernatant using HisTrap FF (GE Healthcare, Uppsala, Sweden). The following buffers were used: wash/binding: 50 mM Tris (pH 10), 300 mM NaCl, and 10 mM imidazole; elution buffer: 50 mM Tris pH 10, 300 mM NaCl, and 500 mM imidazole. Gradient of elution ranging from 0-100 of elution buffer, elution occurred with 25% elution buffer. Fractions containing His6-tagged procruzain, obtained upon elution, were combined, and covalent reversible inhibitor MMTS, final concentration of 2 mM, was added. Eluates were concentrated using Amicon Ultra centrifugal filter units (10 kDa molecular weight cutoff, Millipore). The buffer was changed to 0.1 M acetate (pH 5.5), 0.9 M NaCl, and 10 mM EDTA using NAP-5 desalting column (GE Healthcare).

**Zymogen procruzain auto-activation.** For procruzain auto-activation, the pH was adjusted to 5.3 and 5 mM dithiothreitol (DTT) was added to remove the methylthio protection from Cys25. The reaction time for processing procruzain into active cruzain was 20−45 min at 37 °C. The reaction was monitored visually by observing opacity of the protein solution, which became clear once reaction was complete. Transformation of the inactive zymogen (∼37 kDa) into the catalytically active domain (cruzain, ∼23 kDa) was confirmed by removing aliquots of the solution at selected time points. The reaction was terminated by placement of the sample tubes on ice. These samples were analyzed by sodium dodecyl sulfate−polyacrylamide gel electrophoresis under denaturing conditions and by a fluorescence-based activity assay using the peptide substrate Z-Phe-Arg-AMC [48]. After activation, DTT was removed by desalting on the NAP-5 column (GE Healthcare) in the activation buffer free of the reducing agent. Then cruzain was immediately inhibited with Gallinamide A or MMTS at 1 mM to prevent self-degradation. A fluorescence-based activity assay confirmed enzyme inhibition.

**Cruzain purification**. The last step of purification was performed using the size-exclusion FPLC on the Superdex 75 10/300 GL (GE Healthcare) column. The buffer used in this purification step did not contain reducing agents. To keep cruzain inhibited, MMTS or gallinamide A was added to each of the fractions immediately upon elution from the column. Protein samples were concentrated before and after size-exclusion chromatography to a concentration of approximately 10 mg/ml using Amicon Ultra centrifugal filter unit (3 kDa molecular weight cutoff, Millipore). Protein concentration was assayed by UV-vis spectroscopy, using the molar extinction coefficient of $\varepsilon^{280}$= 68,910 $M^{-1}cm^{-1}$ and 60,430 $M^{-1}cm^{-1}$ for His6-GS-procruzain and cruzain, respectively.
